# An unbiased drug screen for seizure suppressors in Dup15q syndrome reveals 5HT_1A_ and dopamine pathway activation as potential therapies

**DOI:** 10.1101/2020.02.18.954560

**Authors:** Bidisha Roy, Jungsoo Han, Kevin A. Hope, Tracy L. Peters, Glen Palmer, Lawrence T. Reiter

**Author notes:** Corresponding Author: Lawrence T. Reiter, Ph.D., Department of Neurology, University of Tennessee Health Science Center, 855 Monroe Ave, Link 415, Memphis, TN 38103.

## Abstract

Duplication 15q syndrome (Dup15q) is a rare neurogenetic disorder characterized by autistic features and difficult to control (pharmacoresistant) epileptic seizures. Most individuals with isodicentric (idic15) have been on multiple medications to control their seizures and some are still seizing after years of treatment. We recently developed a model of Dup15q in Drosophila by elevating levels of fly Dube3a in glial cells, not neurons. Unlike other Dup15q models, these flies develop seizures that worsen as flies age. Here we used this new model to screen for previously approved compounds from the Prestwick Chemical Library which are able to suppress seizures in flies over-expressing Dube3a in glia using the pan glial driver *repo*-GAL4. We identified 17 out of 1280 compounds in the library that could suppress a bang sensitive (seizure) phenotype. Eight of these compounds were able to suppress seizures significantly in both males and females by at least 50%. Half of these strong seizure suppressors regulated either serotoninergic or dopaminergic signaling and subsequent experiments confirmed that seizure suppression occurs through stimulation of serotonin receptor 5-HT_1A_ but can be further suppressed with the addition of L-Dopa (Levodopa). We provide further support for a seizure model where Dube3a regulation of the Na+/K+ exchanger ATPα in glia can also be modulated by serotonin/dopamine signaling. Finally, based on these pharmacological and genetic studies, we present an argument for the use of 5-HT_1A_ agonists in the treatment of Dup15q epilepsy.

## Introduction

Chromosomal deletions and duplications of the 15q11.2-q13.1 region cause some of the most devastating neurogenetic syndromes in humans, including Angelman syndrome (AS), Prader-Willi syndrome (PWS) and one of the most common genetic abnormalities leading to autism spectrum disorder (ASD), Duplication 15q syndrome (Dup15q) (1). In fact, duplications of 15q are often found in clinical cohorts of ASD, mostly isodicentric duplications of 15q which result in four copies of the genomic region containing the imprinted gene *UBE3A* (2). Multiple lines of evidence from both human genetic studies and experimental models suggest a tight regulation of *UBE3A* expression in the mammalian brain, resulting in maternal allele specific expression of *UBE3A* in neurons of the brain (3–5). In AS, loss of the maternal *UBE3A* allele through 15q11.2-q13.1 deletions, mutations of maternal *UBE3A* or paternal uniparental disomy for the locus all result in an AS phenotype (6). While some evidence suggests that maternal, but not paternal, duplications of 15q result in ASD phenotypes, implicating the maternal expression of UBE3A in neurons (7, 8), there is still not definitive proof from murine models that this is the case (9). Furthermore, multiple studies focused on maternal expression of *Ube3a* in mouse neurons have failed to recapitulate the common and severe pharmacoresistant epilepsy phenotype shared by most isodicentric Dup15q individuals (9–12).

A common co-incident phenotype in ASD is an increased incidence of epileptic seizures (13–15). Although the most prominent feature of Dup15q syndrome is ASD and behavioral abnormalities, most individuals with idic15 also suffer from difficult to control seizures (pharmacoresistant) (1, 16). Typically individuals with idic15 have been on several medications to control their seizures and some individuals continue to suffer from epilepsy despite several anti-epileptic drugs available which can control seizures in AS or even epilepsy of unknown genetic origin. Parent reporting indicates that these pharmacoresistant seizures are the most difficult issue to address in the Dup15q population (16).

Our lab recently developed a model of Dup15q syndrome in *Drosophila melanogaster* which recapitulates the seizure phenotype seen in humans (17). We showed that glial expression, but not neuronal expression, of Dube3a (the fly UBE3A orthologue) results in a robust and reproducible seizure phenotype in flies (17). Although there are currently several mouse models of Dup15q syndrome which elevate Ube3a levels in neurons, none of these mouse models show a spontaneous or easy to induce seizure phenotype like our Drosophila model. In fact, flies expressing *Dube3a* or human *UBE3A* in glial cells will show seizure phenotypes after vortexing, at elevated temperatures or even through optical stimulation using a strobe light (17).

Several lines of evidence from both murine models and human brain studies indicate that the *UBE3A* gene escapes allele specific expression in glial cells of the mammalian brain. Early studies of imprinting in cultured neurons or glia from Ube3a maternal allele deficient mice indicated that not only is *Ube3a* expressed from the paternal allele in cultured glia, but that the *Ube3a* antisense transcript, which regulates *Ube3a* expression on the paternal allele, could not be detected in cultured glia from *Ube3a*^*M−/P+*^ animals (18). Additional studies in the mouse brain showed that *Ube3a* escapes allele specific regulation in GFAP+ glial cells and oligodendrocytes of the brain (19, 20). Finally, studies utilizing a *Ube3a*-YFP knock in construct further validated the finding that *Ube3a* is bi-allelically expressed in glial cells (21). Recently, a study using human brain samples confirmed at the protein level that UBE3A is present in GFAP+ glial cells of the human brain (19).

Given the body of evidence supporting biallelic expression of *UBE3A* in glia and the hypothesis that biallelically expressed *UBE3A* in glial cells should be elevated in idic15 individuals who have 4 copies of *UBE3A*, we initiated a screen using our new fly model of Dup15q to identify small molecules that could suppress seizures and reveal mechanisms through which elevated UBE3A in glia could result in a seizure phenotype. Here we exploited a 30 second window of paralysis observed in all flies expressing *Dube3a* in glia to design a screen for small molecules that can suppress seizures in an effort to identify previously approved drugs that can be repurposed to potentially treat seizures in humans with Dup15q.

## Methods and Materials

### Fly Stocks

All flies were maintained at 25°C on a 12-hour light/dark cycle and raised on standard corn meal media (Bloomington Stock Center). The following stocks were obtained from the Bloomington Drosophila Stock Center (Bloomington, IN): Ddc^DE1^, 5HT1A^Δ5kb^/CyO, DAT^Z21744^, Ddc^27^pr1/CyO, Ddc^k02104^, ple^4^/TM3,Sb, Trh^c01440^, 5-HT2A^C1644^/TM3Sb, 10XUAS-IVS-myr::tdTomato and *repo*-GAL4. The UAS*-ATPα* line (RRID:FlyBase_FBst0500173) was obtained from FlyORF (22). The *Dube3a* and human *UBE3A* UAS lines used in this study have been described previously (23). A complete list of stocks used can be found in **Table S2**.

### Chemical Compound Preparation

The Prestwick Library of 1280 approved drugs was purchased from Prestwick Chemicals (Illkirch, France). This library has high chemical and pharmacological diversity with known bioavailability and safety in humans. All compounds were dissolved in DMSO (Sigma-Aldrich; St. Louis, MO) and mixed with standard corn meal medium to make drug food (0.1% final DMSO concentration; Drug Concentration: 1μM) for the primary screening. For the secondary screening and the other pharmacological interventions, drugs were dissolved in water, ethanol or DMSO (based on their highest solubility) and mixed with standard corn meal medium to make drug food (DMSO concentration used: 0.02% in all drug food except for Ceforanide = 0.05%). Drugs were serially diluted to concentrations: 0.04μM, 0.2μM, 1.0μM and 5.0μM depending on the experiment.

### Primary Seizure Susceptibility Screen

For all behavior tests, flies were not exposed to CO_2_ for at least two days prior to testing to avoid issues related to seizure resistance after exposure to CO_2_. The bang sensitivity assay (BSA) was performed as previously described (24–26). Both drug and non-drug (DMSO alone) fed flies were transferred to empty standard fly vials and subjected to mechanical stress (“bang”) by vortexing the vial at full speed on a standard laboratory vortexer (LabNet) for 10 seconds. After vortexing, but within a 30 second critical window, the flies were separated (seizing from non-seizing) using the apparatus described in **Figure 1B**. This apparatus was constructed by placing a funnel over a falcon tube cut at both ends and placed on top of a Petri dish without a lid. The funnel was subsequently removed and replaced with an empty vial. A timer was started and after 30 seconds, the Petri dish containing the immobile seizing flies was covered with the lid. Subsequently, the falcon tube – empty vial was flipped to transfer all the non-seizing flies into the empty vial. Flies of both the genotypes: *repo*>*Dube3a* and UAS*-Dube3a;TM3,Sb* were counted in the seizure group (Petri dish) and the non-seizure group (empty vial over the falcon tube). Since seizure was induced only in the *repo*>*Dube3a* flies, we used these counts in the final analysis of drug induced seizure recovery. For all the experiments in the primary screen we used 3 - 4 day old flies.

**Figure 1.**
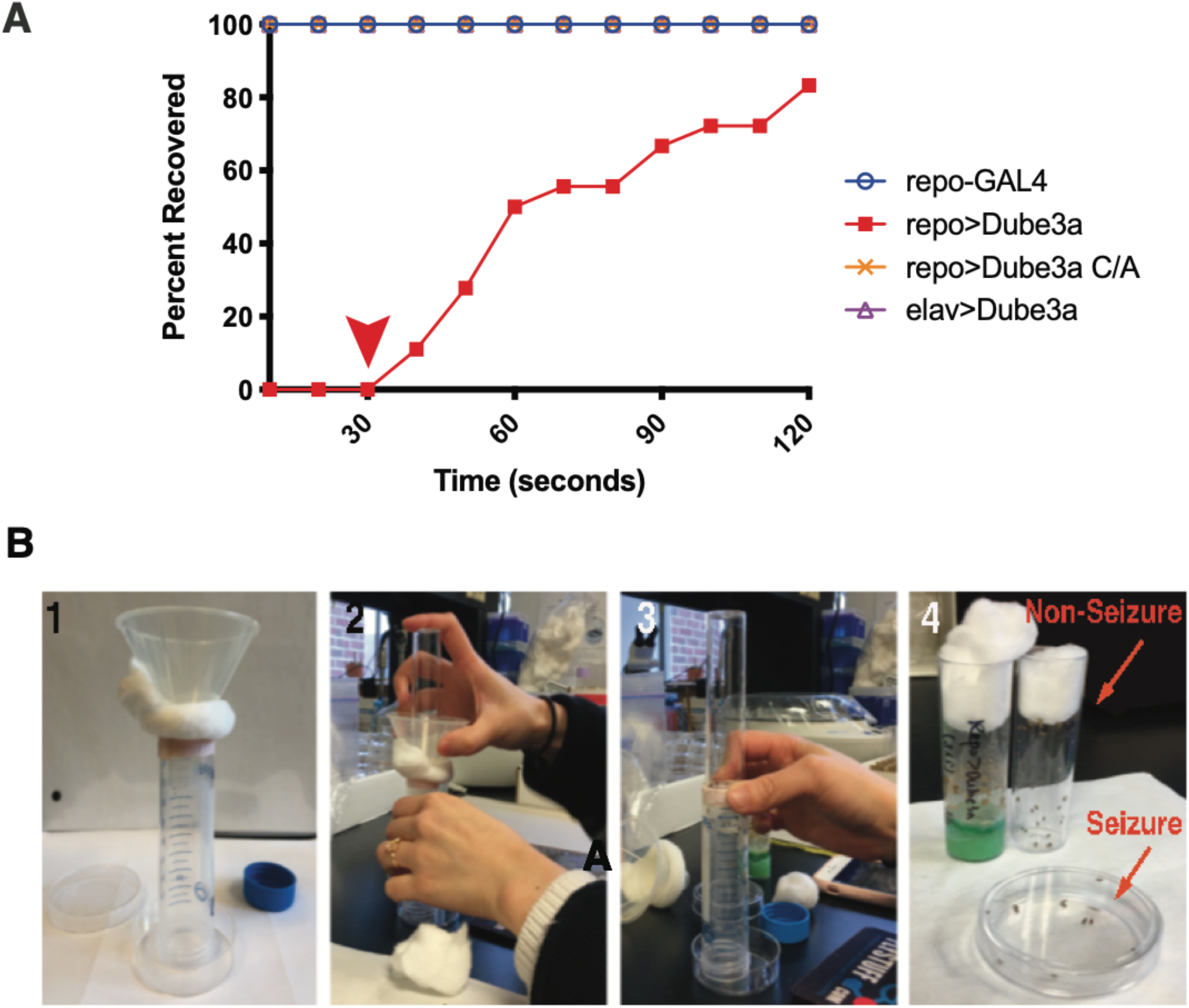
Primary Screening Protocol. A) Only flies of the *repo*>*Dube3a* genotype have bang sensitivity (red filled squares). *elav*>*Dube3a* (purple triangle), *repo*-GAL4 alone (blue open circles) and *repo*>*Dube3a*-C/A animals that are ubiquitin ligase dead (tan cross) all show no bang sensitive phenotype (top line). *repo*>*Dube3a* flies always take at least 30s to begin to recover from seizures (red line and red arrow). Total recovery time for all flies of the *repo*>*Dube3a* genotype is about 80% recovery at 120s. This means there is a window of 30s during the paralysis phase of the seizure when all seizing flies are immobile. B) In panel #1 we show the screening rig consisting of a funnel, a 50ml conical tube cut at the bottom and a petri dish with a lid. In #2 you can see flies that were vortexed for 10s are dumped into the funnel. Flies that are seizing drop into the petri dish, while flies that have recovered can climb up the vile into a new tube (#3). After no more than 30s, the flies in both groups are collected with seizing flies in the dish and non-seizing flies in an empty vial (#4).

### Secondary Validation Seizure Assays

Based on our previous study of glial initiated seizures, we used 4 day old *repo*>*Dube3a* and 9 day old *repo*>*UBE3A* flies (17) in subsequent experiments. For all behavior tests, flies were sorted and separated at least two days prior to testing to avoid complications from CO_2_ anaesthesia. Flies of each genotype, *repo*>*Dube3a* or *repo*>*UBE3A* (both with and without drug) were sorted by sex post eclosion and transferred to freshly prepared drug containing food or solvent-only food.

For seizure behavior studies, flies were anesthetized briefly by exposing the vials to ice for 2-3 minutes. Then 3-5 flies were distributed to empty vials for the bang sensitivity assay. After a 1-hour recovery from cold-anesthesia, the flies were subjected to mechanical stress (“bang”) by vortexing the vial using a standard laboratory vortexer (LabNet) at top speed for 10 seconds. The total recovery time was recorded for each individual fly in the vial using a digital timer. Total recovery time in our assay was defined as the time taken by a fly to fully recover (i.e. when the fly righted itself and was able to walk freely around on the base or on the walls of the vial). For each drug at each concentration, or combination of drugs, at least 40 flies of the correct genotype were tested in the bang sensitivity assay.

### Glial Cell K^+^ Content Assay

Stock solutions of 1 mM Asante Potassium Green 2 (APG-2, TEFlabs), a fluorescent cell membrane permeable K^+^ indicator, were prepared in DMSO. Three-day old *repo*>*tdTomato* or *repo*>*Dube3a*+*tdTomato* flies fed on food with water or food with 0.04μM 5HT + L-dopa were briefly anesthetized with CO_2_ and their heads were removed. Heads were submerged in room temperature Drosophila saline consisting of the following (in mM): 128 NaCl, 1.8 CaCl_2_, 2 KCl, 5 MgCl_2_, 36 sucrose, 5 HEPES, pH 7.2. Brains were dissected from the head case and incubated with 7.5μM APG-2 in Drosophila saline at room temperature with gentle agitation for 1 hour. Brains were then washed 2X with Drosophila saline and wet mounted on a microscope slide with Drosophila saline. Images were captured from the optic lobe on a Leica DM6000B microscope (Leica) using a 63X oil immersion lens using the N3 and L5 filters for tdTomato and APG-2 respectively. Exposure times were calibrated on *repo*>*tdTomato* brains and microscope settings remained constant between *repo*>*tdTomato* and *repo*>*Dube3a*+*tdTomato* groups to allow for direct comparison of fluorescence intensity. APG-2 fluorescence intensity was calculated offline using Adobe Photoshop CC2018 and ImageJ (27). The APG-2 channel was converted to a gray scale image using Adobe Photoshop. Brightness and contrast parameters were kept constant in all images across various groups. Only tdTomato-positive glial cells were selected for analysis. Cells were selected using the “freeform” tool to outline the cell body in the APG-2 channel. Using the “measure” command, the mean fluorescence value for each cell was recorded as previous described (17). A minimum of 35 cells was analyzed per group from at least 3 images each from 3 different brains per group.

### Statistical Analysis and Graphing

All graphs and statistical tests were performed using Prism version 7.0 (GraphPad). All histograms and measurements are shown as mean ± SEM. To test for statistical significance between two samples, Student’s *t*-test with two tailed comparison was used. To determine statistical significance among multiple samples, we used a one-way ANOVA with Tukey’s multiple comparison correction. Statistical significance is indicated on all graphs by convention: p_value_ ≤ 0.05 (*), p_value_ ≤ 0.01 (**), p_value_ ≤ 0.001 (***) and p_value_≤ 0.0001 (****). All analyses were performed with the experimenter blinded to genotype.

## Results

In a previous study, we determined that glial expression of fly *Dube3a* using the pan-glial driver *repo*-GAL4 results in a bang sensitive phenotype, i.e. seizures after vortexing. In addition, we found that pan-neuronal expression using *elav*-GAL4 or expression of a ubiquitin ligase defective form of Dube3a did not evoke seizures (17). These studies also revealed a 30 second window after vortexing wherein 100% of *repo*>*Dube3a* animals were in the paralysis phase (**Figure 1A**). We designed a new screening method based on exploiting this 30 second window of paralysis to separate flies that are not seizing from flies that are seizing. Flies that are seizing will fall to the bottom of the tube after vortexing, but flies that are not seizing, have very short seizure recovery times, or are not the *repo*>*Dube3a* genotype, will crawl towards the top of the tube and can be collected separately (**Figure 1B**). Each group of flies is then evaluated visually under the microscope to identify only *repo*>*Dube3a* flies in each group for scoring. For the drug screen, we used the Prestwick Chemical Library (Prestwick Chemical) of 1280 off-patent small molecules previously approved for use in humans. Food was prepared containing 1μM of each compound plus DMSO only controls. Flies were grown at 25°C on drug containing food from embryos and tested in the modified bang sensitivity screen 3-5 days after eclosion.

### Primary and Secondary Screening and Validation Assays Reveals 8 Candidate Compounds for Seizure Suppression

Our primary screening selection criteria for compounds that could suppress seizure activity was that compounds must suppress seizures in at least 25% of the *repo*>*Dube3a* animals within the 30s collection window. Using these criteria, we identified 17 compounds that could suppress seizures in *repo*>*Dube3a* animals that eclosed from triplicate crosses (**Table 1**). We immediately noted that 5 out of 17 compounds, which passed primary screening, were related in some way to serotonin (serotonin hydrochloride (5HT), mirtazapine and minaprine dihydrochloride) or dopamine (minaprine dihydrochloride, levodopa (L-dopa) and prenylamine lactate) signaling.

**Table 1.** Primary Screen Hits. Percent suppression is the percentage of flies expressing Dube3a in glia that were not seizing. *Only one representative DMSO vial is shown here. For more extensive DMOS data in males and females see Figure S1.

|  |  | <u>Seizure</u> |  | <u>No seizure</u> |  |  |  |
| --- | --- | --- | --- | --- | --- | --- | --- |
| Code | Compound | *repo>Dube3a* | UAS-*Dube3a*; TM3, *Sb* | *repo>Dube3a* | UAS-*Dube3a*; TM3, *Sb* | Total # *repo>Dube3a* | Percent Suppression |
| N/A | DMSO* | 6 | 1 | 0 | 15 | 6 | **0.0%** |
| Prestw-1 | Azaguanine-8 | 8 | 0 | 6 | 17 | 14 | **42.9%** |
| Prestw-17 | Levodopa | 8 | 0 | 4 | 9 | 12 | **33.3%** |
| Prestw-19 | Captopril | 10 | 0 | 6 | 13 | 16 | **37.5%** |
| Prestw-66 | Minaprine dihydrochloride | 1 | 2 | 1 | 1 | 2 | **50.0%** |
| Prestw-390 | Fusidic acid sodium salt | 9 | 0 | 7 | 20 | 16 | **43.8%** |
| Prestw-457 | Meclozine dihydrochloride | 9 | 0 | 7 | 16 | 16 | **43.8%** |
| Prestw-470 | Ceforanide | 21 | 0 | 7 | 23 | 28 | **25.0%** |
| Prestw-1358 | Vatalanib | 21 | 1 | 8 | 47 | 29 | **27.6%** |
| Prestw-1385 | Closantel | 17 | 1 | 7 | 43 | 24 | **29.2%** |
| Prestw-481 | Serotonin hydrochloride | 14 | 0 | 7 | 32 | 21 | **33.3%** |
| Prestw-1144 | Mirtazapine | 10 | 0 | 12 | 19 | 22 | **54.5%** |
| Prestw-560 | Prenylamine lactate | 7 | 0 | 4 | 8 | 11 | **36.4%** |
| Prestw-871 | Iopamidol | 12 | 1 | 7 | 43 | 19 | **36.8%** |
| Prestw-1116 | Dorzolamide hydrochloride | 2 | 2 | 2 | 2 | 4 | **50.0%** |
| Prestw-1118 | Cefepime hydrochloride | 19 | 2 | 7 | 41 | 26 | **26.9%** |
| Prestw-1711 | Felbamate | 3 | 2 | 3 | 6 | 6 | **50.0%** |
| Prestw-1318 | Pranoprofen | 12 | 0 | 8 | 37 | 20 | **40.0%** |

Validation of the 17 primary screen compounds was performed using a more accurate bang sensitivity assay with individual flies and measuring the actual seizure recovery times. All 17 compounds were also tested at 5μM, 1μM, 0.2μM and 0.04μM concentrations in order to identify the lowest concentration that was still effective at seizure suppression and in the appropriate solvent (DMSO, water or EtOH depending on the compound). While there were no differences in seizure activity across these different solvents, we did note a difference in variance in male flies versus female *repo*>*Dube3a* flies (**Figure S1**). For this reason, we tested both males and females separately for the validation assays.

The go/no go criteria for validation of a primary screen candidate was as follows: the compound must suppress seizures by at least a 50% decrease in recovery time versus solvent alone at one of the tested concentrations; 2) the suppression must reach a statistical significance by one-way ANOVA of at least p_value≤_0.01; 3) the suppression must occur in both males and females. Using these criteria we were able to identify 8/17 primary screen candidates that can suppress seizures in *repo*>*Dube3a* males and females (**Figure 2**). Using these stringent criteria we found that most of the compounds could suppress seizures by 50% in males and females at a concentration of 0.04μM in the food (the exception being Iopamidol, which required 1μM and both levodopa and dorzolamide hydrochloride which both required at least 0.2μM for suppression). A complete list of these compounds and their approved uses in humans can be found in **Table S1**. Drugs that failed the secondary validation can be found in **Figure S2**.

**Figure 2.**
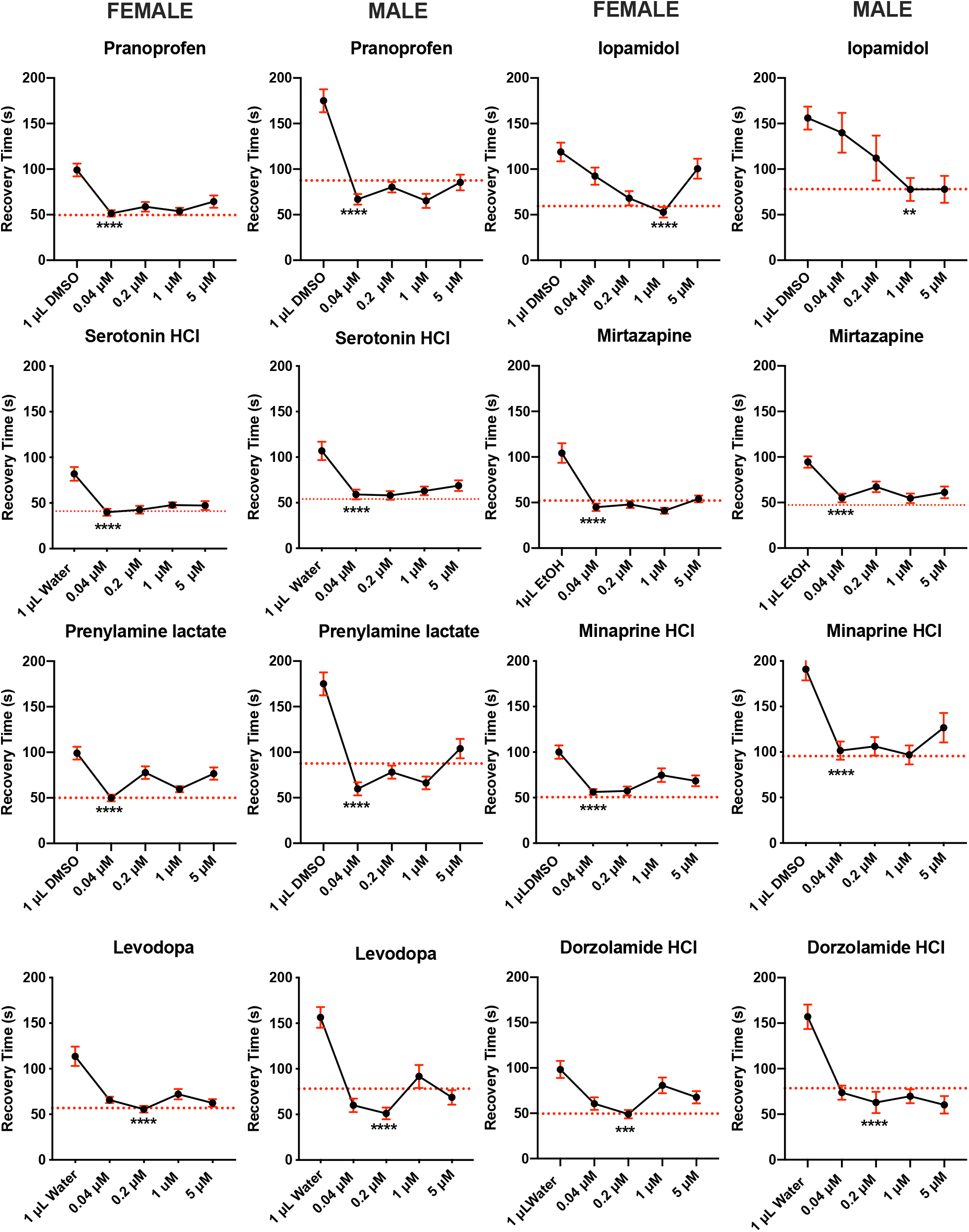
Drugs that Passed Secondary Screening Validation. All 17 compounds that were identified in the primary screen were tested again using a modified single fly bang sensitivity test that measures recovery time after vortexing. Eight compounds were able to suppress seizures at concentrations lower than the primary screen (1μM) and as low as 0.04μM for some compounds. All compounds suppressed seizures by ≥50% of solvent alone (DMSO or Water) in males and female, at p_value_≤0.01. Four of the 8 compounds are associated with serotonin and/or dopamine signaling (serotonin HCl, mirtazapine, minaprine HCl and Levodopa). Error bars (red) are SEM. N>40 animals per data point.

Once again, the five compounds involved in serotonin and dopamine signaling identified in the primary screen which were able to reach significance included 5HT, mirtazapine, minaprine dihydrochloride, prenylamine lactate and L-Dopa. The relationship among the three serotonin signaling compounds provided clues to the mechanism of action of seizure suppression. Serotonin can stimulate 5-HT_1_, 5-HT_2_, 5-HT_3_ and 5-HT_4,5,6,7_ receptors, but mirtazapine is a strong antagonist of all serotonin receptors except 5-HT_1A_ (28–31). Minaprine dihydrochloride is a serotonin and dopamine re-uptake inhibitor, and can therefore increase the levels of free serotonin and dopamine at the synapse (32). Another drug, prenylamine lactate, is known to prevent noradrenaline and dopamine from reuptake by storage granules, thereby increasing their concentration in the synaptic cleft (33, 34). Thus, both minaprine dihydrochloride and prenylamine lactate could possibly increase serotonin and dopamine levels at the synapse, which is consistent with the observation that levodopa can also suppress seizures in addition to serotonin (**Figure 2**). Additional information about these drugs can be obtained from **Ch**emical **E**ntities of **B**iological **I**nterest (ChEBI, EMBL-EBI) and DrugBank databases.

### Combining Compounds with Serotonin Reveals an additive effect of Dopamine Signaling in Suppressing Seizures

Next, we conducted experiments on the combined effect of 5HT with drugs modulating serotoninergic / dopaminergic signaling and other drugs that suppressed seizures in our screen. Three drugs related to serotonin or dopamine signaling (Minaprine HCl, Prenylamine lactate and Mirtazapine) and one unrelated drug (Dorzolamide HCl) were able to significantly decrease recovery time more than serotonin hydrochloride alone in female *repo*>*Dube3a* flies (**Figure S3**). The addition of minaprine hydrochrloride (p_value_≤0.05), prenylamine lactate (p_value_≤0.01), mirtazapine (p_value_≤0.05) plus 5HT could decrease recovery time below the addition of 5HT alone in *repo*>*Dube3a* females, but had little effect on seizure suppression in males. However, the combination of 5HT plus L-dopa did significantly decrease recovery time below 5HT alone in both male and female *repo*>*Dube3a* flies (p_value_≤0.001) (**Figure 3A**). These results show that a combination of serotonin and dopamine signaling may be required for the maximum seizure suppression in flies with elevated Dube3a levels in glia.

**Figure 3.**
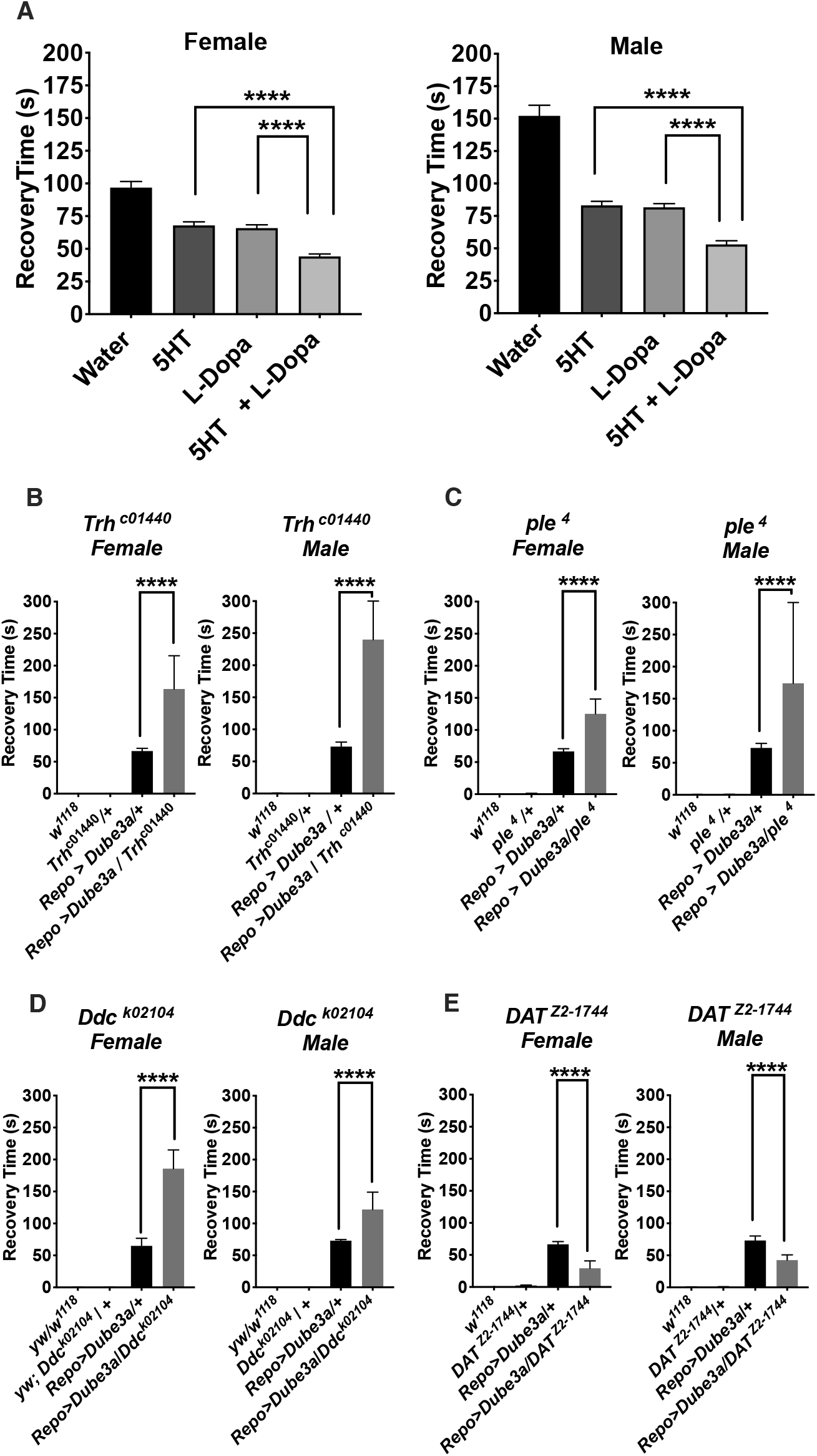
Combining serotonin with dopamine can further suppress seizures. A) Synergistic effects of drugs at four diminishing concentrations indicated that of the 8 compounds identified, only L-dopa could further suppress the effects of serotonin alone in both males and females. Although some combinations were able to decrease recovery times in males or females, no other combination reached significance in both (**Figure S3**). B-D) Mutations in critical enzymes for either serotonin synthesis (*Trh*^*c1440*^), dopamine synthesis (*ple*^*4*^) or shared by both pathways (*Ddc*^*DEI*^) significantly enhanced seizure recovery time in *repo*>*Dube3a* flies. E) Loss of function mutations in the dopamine transporter gene *DAT*^*Z2-1744*^, which clears dopamine from the synaptic space, significantly suppressed seizures in *repo*>*Dube3a* animals, decreasing recovery time. Error bars are SEM. N>100 animals per column for (A), N>7 animals per column for (B-E).

In order to support this pharmacological data with genetic evidence, we acquired several mutants in various components of the serotonin and dopamine synthesis pathways and one mutant involved in dopamine re-uptake, available from the Bloomington Stock Center. The synthesis of both serotonin and dopamine is linked in both flies and humans via an enzymatic pathway that contains three enzymes. Tryptophan hydroxylase (*Trh* in flies) is the first enzymatic step in the conversion of tryptophan to serotonin followed by dopa decarboxylase (*Ddc* in flies) also known as aromatic L-amino acid decarboxylase which coverts 5-hydroxytryptophan to serotonin. Dopamine is made by converting tyrosine to L-dopa via tyrosine hydroxylase (*ple* in flies) and then L-dopa is converted to dopamine via the same L-amino acid decarboxylase as serotonin (*Ddc*). Heterozygous loss of function mutations in *Trh*^*c1440*^, *Dd*^*K02104I*^ or *ple*^*4*^ significantly increased seizure recovery time, by as much as 2X, over *repo*>*Dube3a* animals alone (**Figure 3B-D**), indicating that down regulation of serotonin or dopamine exacerbates the seizure phenotype. In addition, a heterozygous loss of function mutation called *DAT*^*Z2-1744*^ in the dopamine transporter gene (*dopamine active transport* or *DAT* in flies) which clears dopamine from the synaptic space, significantly suppressed seizures in *repo*>*Dube3a* animals (**Figure 3C**). These data suggest that more dopamine in the synaptic space has a therapeutic effect on seizures produced by elevated Dube3a in glia. In combination with the neurotransmitter synthesis data, these genetic studies fully support the pharmacological studies which suggest stimulation of serotonin and/or dopamine pathways can significantly suppress seizures in our model.

### Specific Stimulation of 5-HT_1A_ Receptor Alone Suppresses Seizures in Repo>Dube3a Flies

Since both serotonin, which stimulates all serotonin receptors, and mirtazapine, which is an antagonist of all serotonin receptors except 5-HT_1A_, were both able to suppress seizures in *repo*>*Dube3a* flies, we decided to broaden our analysis using a series of specific serotonin receptor agonists and antagonists for their ability to suppress or enhance seizures in order to address the hypothesis that stimulation of 5-HT_1A_ alone can suppress seizures. **Table 2** lists specific 5-HT receptor agonists and antagonists, as well as the predicted result if we feed *repo*>*Dube3a* animals these compounds and test for seizure activity. Two 5-HT_1A_ agonists (serotonin hydrochloride and U92016A), two 5-HT_1A_ antagonists (WAY100635 and mianserin HCl), one 5-HT_2A_ antagonist (ketanserin tartrate) and one agonist for 5-HT_2A,2B,2C_, but not 5-HT_1_ (https://www.wikidata.org), were assayed for their ability to suppress or enhance seizures at concentrations from 0.04μM to 5μM. All drugs that stimulated 5-HT_1A_, directly or indirectly, were able to suppress seizures at concentrations down to 0.04μM in the food (**Figure 4A**). All compounds that either directly or indirectly routed serotonin away from 5-HT_1A_ increased the seizure recovery times significantly in *repo*>*Dube3a* animals at concentrations ranging from 0.04μM to 1μM in the food (**Figure 4B**). In fact, these same agonists and antagonists had identical effects on seizures in 9 day old flies expressing human UBE3A in glia at identical dosages that were effective in animals expressing fly *Dube3a* in glia (**Figure 5**). Together, these results indicate that stimulation of 5-HT_1A_, or inhibition of 5-HT_2A_ in the presence of active 5-HT_1A_, can pharmacologically suppress seizures generated in glia by over-expression of either fly or human *UBE3A*.

**Table 2.** Serotonin Receptor Agonists/Antagonists and Predicted Effects on Seizure.

| Drug | Action | Specificity | Seizure Prediction |
| --- | --- | --- | --- |
| Serotonin HCl | Agonist | All 5-HT Receptors | *Suppressor* |
|  |  | 5-HT 1,2,3,4,5,6,7 |  |
| Mirtazapine | Antagonist | 5-HT2A, 5-HT2C, 5-HT3 | *Suppressor* |
| (R) DOI HCl | Agonist | 5-HT2A, 5-HT2B, 5-HT2C | **Enhancer** |
| Ketanserin tartate | Antagonist | 5-HT2A | *Suppressor* |
| WAY100635 | Antagonist | 5-HT1A | **Enhancer** |
| Mianserin HCl | Antagonist | All 5-HT Receptors | **Enhancer** |
|  |  | 5-HT 1D, 1F, 2A, 2B, 2C, 3, 6, 7 |  |
| U92016A | Agonist | 5-HT1A | *Suppressor* |

**Figure 4.**
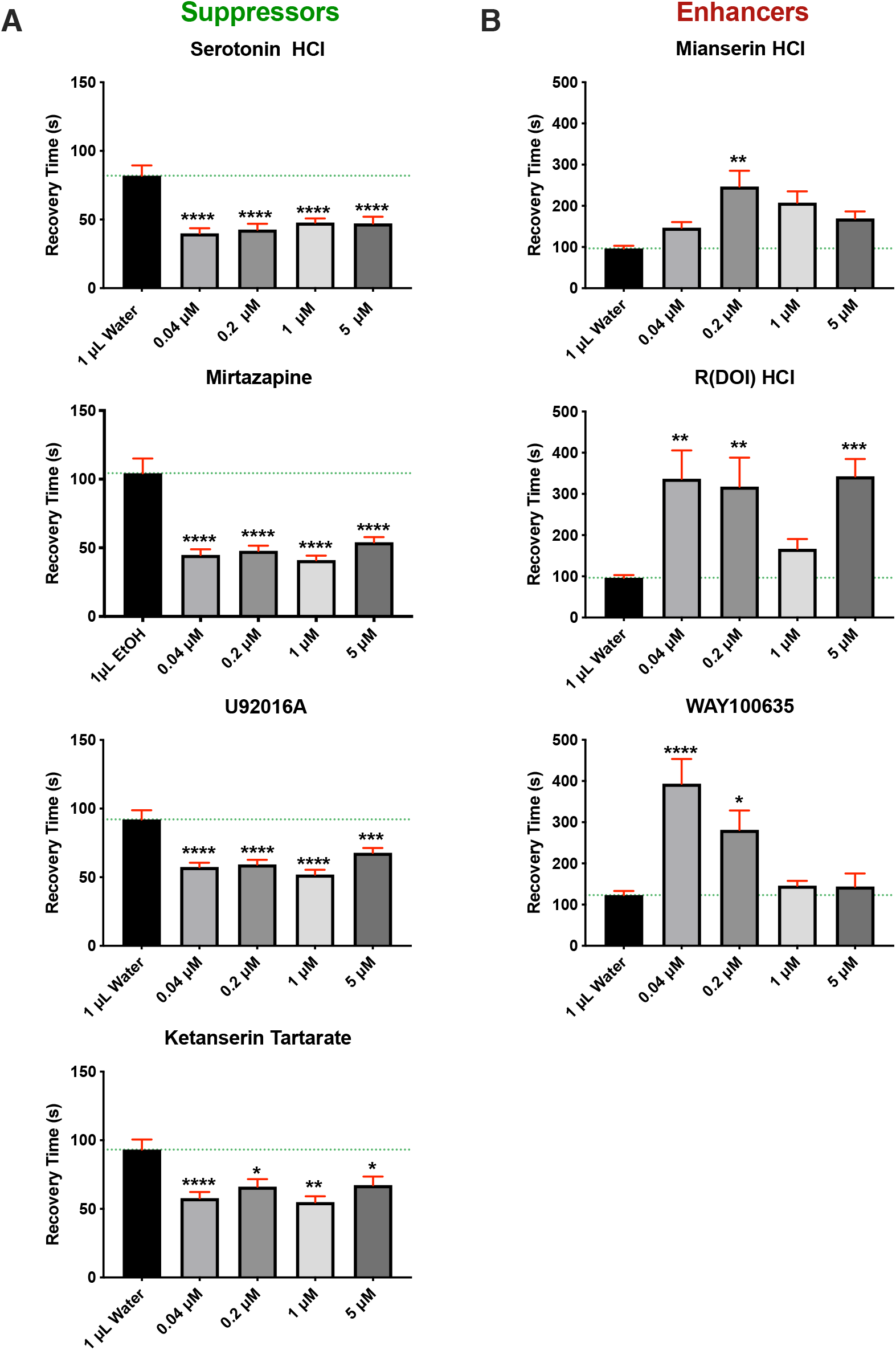
Serotonin Agonists and Antagonists Reveal a 5-HT_1A_ Mechanism for Suppression. Flies expressing *repo*>*Dube3a* were assayed for bang sensitivity (total recovery time in seconds) after being raised on food containing solvent alone (water or EtOH) versus increasing concentrations of various 5-HT_1A_ agonists and antagonists. A) All compounds that act either as 5-HT_1A_ agonists (Serotonin HCl, U92016A and Ketanserin Tartrate) or are antagonists to 5-HT_2A_ receptor (Mirtazapine) significantly suppressed seizures in *repo*>*Dube3a* animals at concentrations down to 0.04μM in the food. B) All compounds that are 5-HT_1A_ antagonists (Mianserin HCl, R(DOI) HCl, WAY100635) significantly enhanced seizure recovery time at concentrations between 0.04-0.2μM. Green lines are average recovery time in water or EtOH. Error bars (red) are SEM. N>60 animals per data point. All flies in this figure were female, males showed similar results (**Figure S4**).

**Figure 5.**
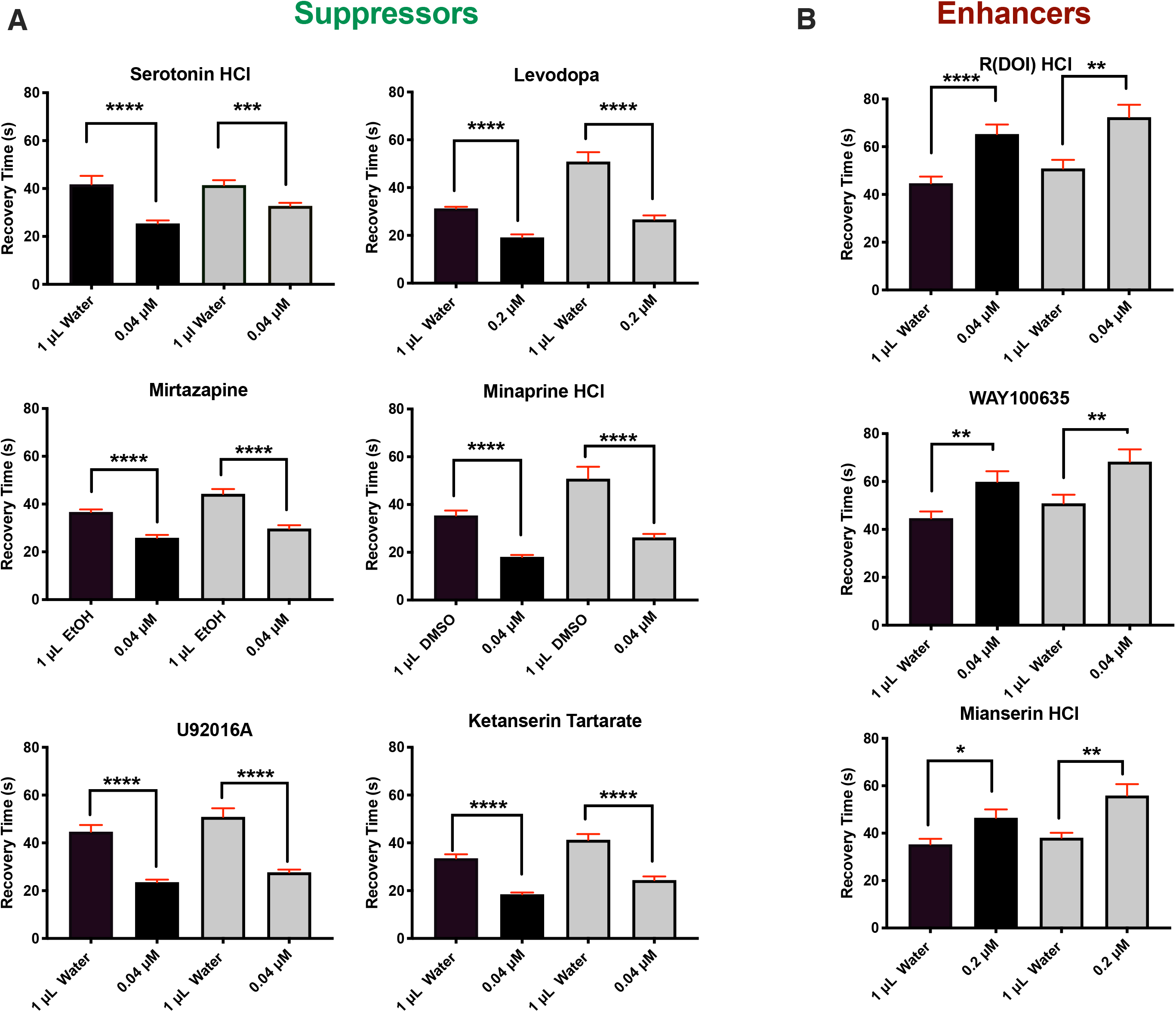
Seizures in *repo*>*UBE3A* Animals also Respond to 5-HT_1A_ Agonists. As with flies expressing fly Dube3a, flies expressing human UBE3A in glial cells also show decreased seizure recovery time when treated with 5-HT_1A_ agonists (A), but increased seizure recovery times when treated with 5-HT_1A_ antagonists (B). Females are black columns and males are grey. Error bars (red) are SEM. N>70 animals per data point.

In order to support these pharmacological findings, we performed genetic experiments designed to determine if serotonin could still suppress seizures in *repo*>*Dube3a* flies in a 5-HT_1A_ or 5-HT_2A_ mutant background. Loss of function alleles of both 5-HT_1A_(*5HT1A^Δ5kb^*) or 5-HT_2A_(*5-HT2A^C1644^*) receptors were identified in the BDSC which were then crossed to a stock containing both *repo*-GAL4 and UAS-*Dube3a* over X*ap* (*w*^*1118*^; [*repo*-GAL4;UAS-*Dube3a*]/X*ap*). As expected, loss of 5-HT_1A_ significantly increased seizure recovery time in *repo*>*Dube3a* animals, but loss of 5-HT_2A_ had no effect (**Figure 6A)**. There was also a noticeable shift in transcript expression from 5-HT_1A_ to 5-HT_2A_ in *repo*>*Dube3a* flies vs control animals (**Figure 6B**). Although the pharmacological data indicated that inhibition of 5-HT_2A_ can also suppress seizures, these genetic experiments show that the 5-HT_2A_ receptor itself is not involved in seizure suppression, suggesting that inhibition of 5-HT_2A_ pharmacologically simply frees up more serotonin in the synaptic space which can then stimulate the 5-HT_1A_ to suppress seizures.

**Figure 6.**
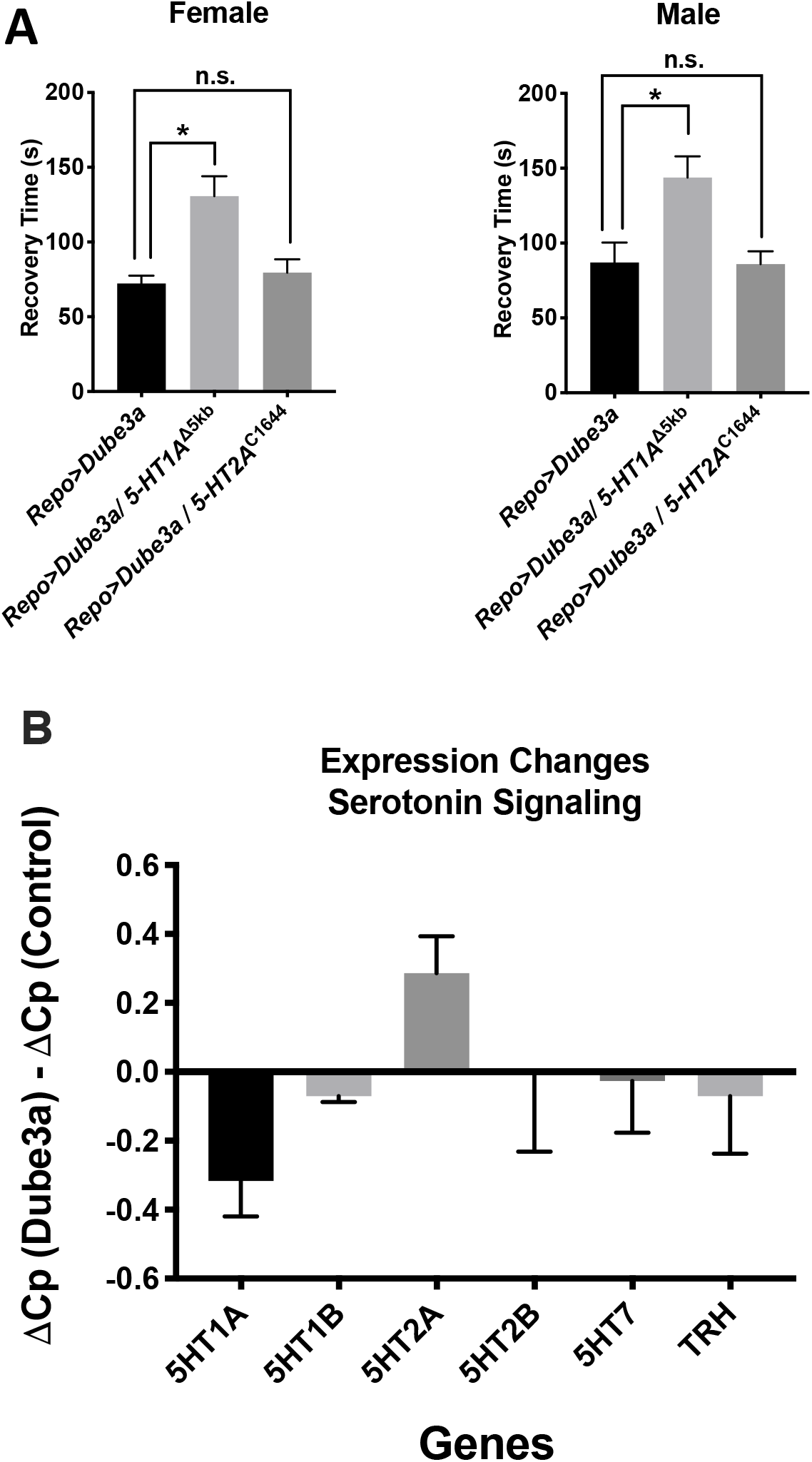
Mutations in 5-HT_2A_ Have no Effect on Seizure Suppression by Serotonin. A) While mutations in 5-HT_1A_ alone do not show a bang sensitive phenotype in *repo*>*Dube3a* animals, the addition of a 5-HT_1A_ loss of function mutation significantly enhances the seizure recovery time even in the presence of serotonin in the food. Loss of function mutations in 5-HT_2A_ showed no bang sensitive phenotype on their own and did not affect seizure recovery time in *repo*>*Dube3a* flies. Error bars are SEM. N>18 animals per data point. B) qRT-PCR gene expression data indicates that the transcription levels of *5-HT*_*1A*_ decrease and the transcription of *5-HT*_*2A*_ increases in the brains of *repo*>*Dube3a* animals relative to *repo*-GAL4 control brains.

In a previous study, we showed that glial overexpression of *Dube3a* causes seizures and synaptic impairments in Drosophila concomitant with down regulation of the Na^+^/K^+^ pump ATPα (17). Serotonin and dopamine are known activators of glial or astrocytic Na^+^/K^+^ pump ATPα via adenyl cyclase dependent protein kinase A (PKA) signaling (35–37). In order to understand the mechanism underlying serotonin or dopamine mediated seizure suppression in *repo*>*Dube3a* flies, we fed serotonin or dopamine to flies co-expressing *ATPα*, which we previously showed could partially rescue seizures (17). We found that either serotonin or dopamine suppressed seizures more effectively in flies co-expressing *ATPα* and *Dube3a*, than flies expressing either *ATPα* + *Dube3a* (without drug) or *Dube3a* (with drug) (**Figure 7A-B**). Finally, adding a cocktail of both serotonin and dopamine while co-expressing *ATPα* almost completely suppressed seizures in *repo*>*Dube3a* flies (**Figure 7C**).

**Figure 7.**
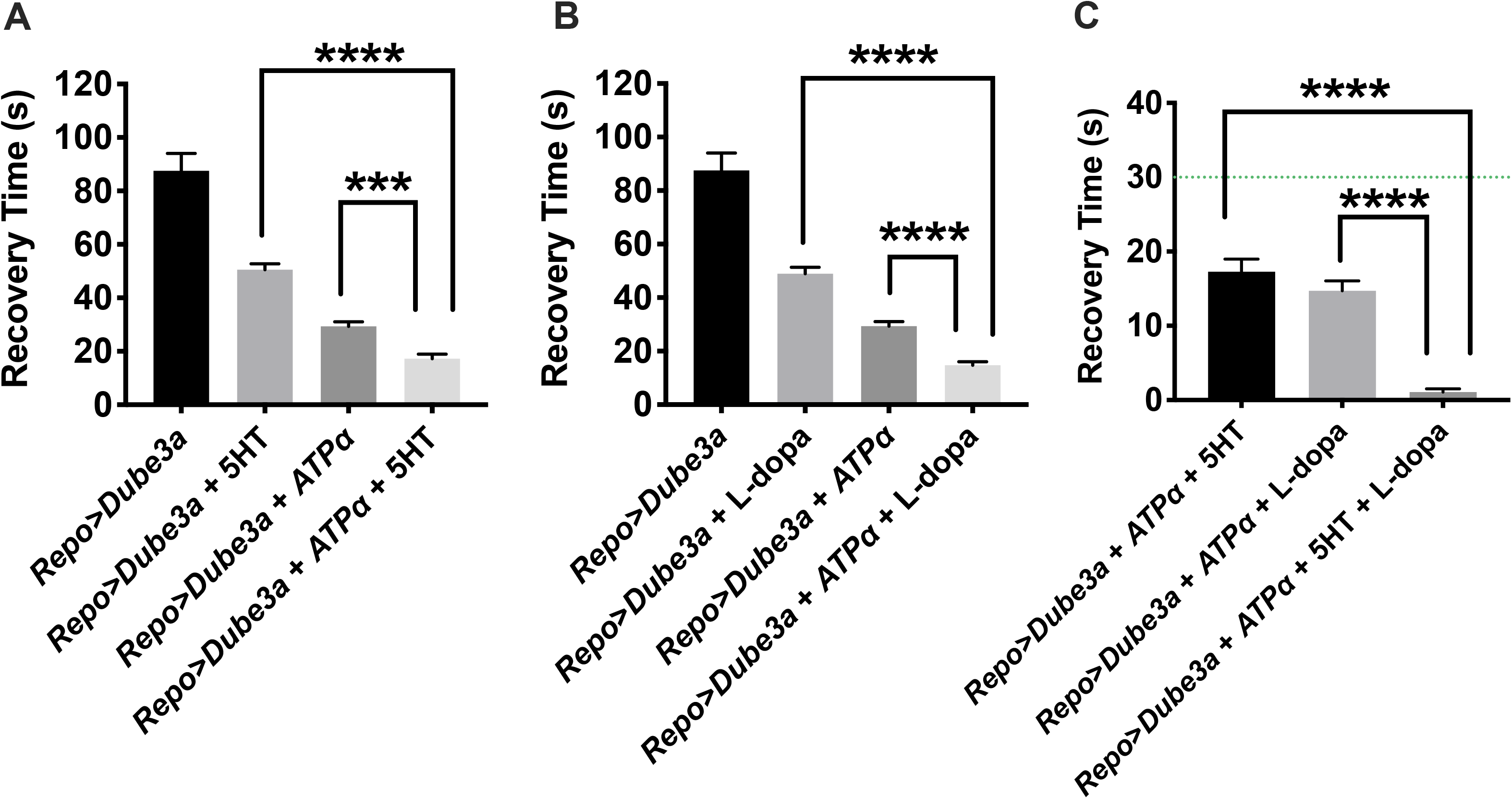
Complete rescue of seizure phenotype in *repo*>*Dube3a* animals. A) Expression of ATPα with 5HT treatment significantly decreases recovery time from >85s to <20s on average. B) Expression of ATPα with L-dopa treatment significantly decreases recovery time from >85s to <20s on average. C) Expression of ATPα with 5HT + L-dopa treatment significantly decreases recovery time from >80s to <1s on average. Most animals had no seizures at all. The green line is 30s, the minimum recover time for all *repo*>*Dube3a* flies we used for primary screening. Error bars are SEM. N>20 animals per column.

The function of ATPα is to pump K^+^ into the cell while pumping Na^+^ into the extracellular space in an ATP-dependent manner. Previously, we showed that elevated Dube3a levels within glia reduce internal K^+^, consistent with the hypothesis that Dube3a down regulates ATPα levels causing a disruption of Na^+^/K^+^ homeostasis (17). We dissected whole brains from *repo*>*tdTomato*, *repo*>*tdTomato* + *Dube3a*,*repo*>*tdTomato* + *Dube3a* (fed 5HT) and *repo*>*tdTomato* + *Dube3a* (fed L-dopa) animals in order to visualize K^+^ concentrations within glia. Brains were incubated in a cell-permeable fluorescent K^+^ indicator called Asante Potassium Green 2 AM (APG-2) as in our prior study (17). Quantification of the fluorescence intensity of APG-2 in tdTomato^+^ cells is an indicator of the relative concentration of K^+^ in glial cells across all genotypes. As expected, there was a significant reduction in APG-2 fluorescence in *repo*>*tdTomato* + *Dube3a* compared to control *repo*>*tdTomato* glial cells (**Figure 8**). However, the addition of a serotonin + L-dopa cocktail restored the concentration of K^+^ within the tdTomato^+^ glial cells (**Figure 8C**). These data suggest that 5HT and L-dopa can modulate ATPα activity in glia, restoring intercellular K^+^ levels, and thereby maintaining the ionic homeostasis at the synapse.

**Figure 8.**
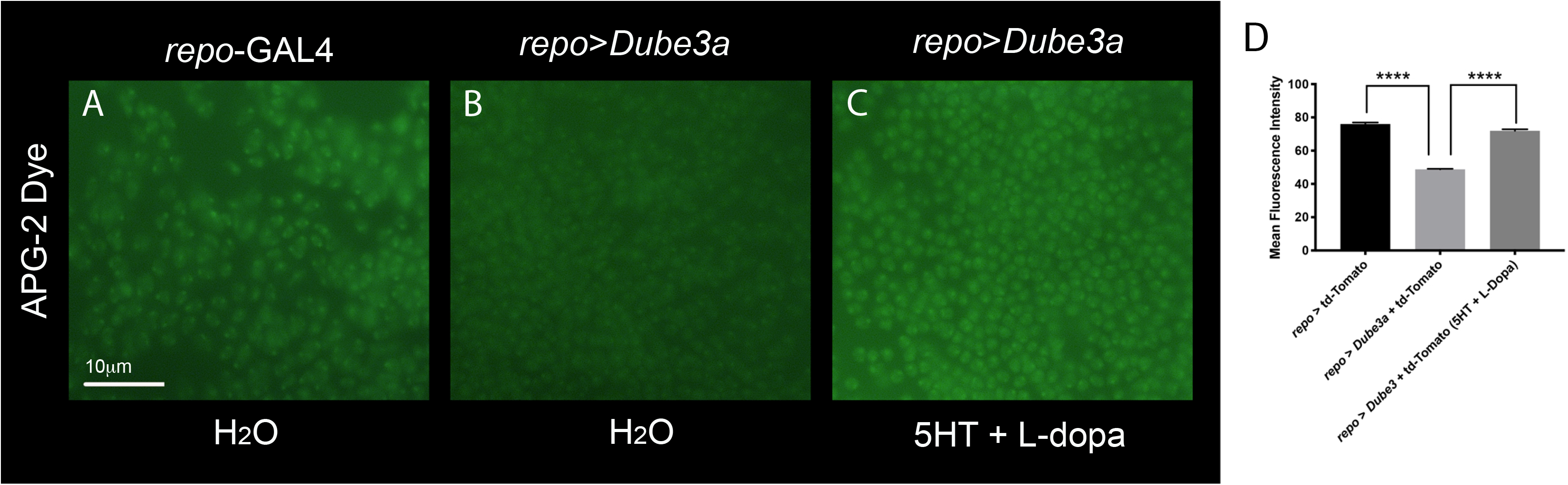
Restoration of K^+^ levels in glial cells though pharmacological intervention with 5HT and L-dopa. Fresh dissected fly brains incubated in the K^+^ binding fluorescent dye APG-2 (green) visualized at 100X on an upright fluorescent microscope. All images were taken using the same exposure settings. A) *repo-*GAL4 alone shows a moderate level of K^+^ within glial cells also marked by *repo*>*td-tomato* (red channel not shown). B) *repo*>*Dube3a* brains show a significant decrease in the amount of K^+^ detected in glia. C) *repo*>*Dube3a* animals raised on food containing a cocktail of 5HT + L-dopa show complete restoration of K^+^ levels in glia. D) Quantification of mean fluorescent intensity using 63X images analyzed as described in the methods section using ImageJ. APG-2 signal quantification confirmed a significant rescue of K^+^ levels in glia for *repo*>*Dube3a* flies when raised on 5HT + L-Dopa. Error bars are SEM. N>35 cells per column.

## Discussion

### A Role for Serotonin Signaling in Dup15q Seizure Suppression

Epilepsy is a common co-morbidity with ASD in general (15, 38–43), but can be even more severe and difficult to treat in syndromic forms of ASD like Dup15q syndrome (1, 16, 44, 45). In fact, most children with isodicentric duplications of 15q that include four copies of the *UBE3A* gene, presumably active in glia where it is not imprinted (18–21), suffer from pharmacoresistant seizures(16). Here we utilized our fly model of Dup15q to screen for previously approved compounds that can suppress seizures due to elevated levels of UBE3A in glia, not neurons (17). We identified 8 compounds that could significantly suppress these gliopathic seizures in both males and females. Surprisingly, five of these compounds (serotonin, levodopa, mirtazapine, prenylamine and minaprine dihydrochloride) are specifically related to serotonin and dopamine signaling. We show that this signaling is preferentially through stimulation of serotonin 5-HT_1A_ receptors, although we are still not sure which specific dopamine receptors may be involved in seizure suppression. The pharmacological and genetic data presented here strongly suggest that specific agonists of 5-HT_1A_ used to treat depression, for example the drug vortioxetine, could potentially be strong suppressors of seizure activity in Dup15q individuals.

### A Possible Mechanism for Seizure Suppression

We found that inhibition of or heterozygous mutations in 5-HT_2A_ alone do not influence seizure susceptibility in *repo*>*Dube3a* flies (**Figure 6A**). One explanation is that in the presence of 5-HT_2A_ antagonists or background mutations in 5-HT_2A_ receptor, free serotonin is still able to suppress seizures in *repo*>*Dube3a* flies through 5-HT_1A_. Cross talk between 5-HT_1A_ and 5-HT_2A_ receptors has been documented in the mammalian brain previously (46). Our own transcript expression studies here indicate that overexpression of Dube3a simultaneously down regulates 5HT_1A_ receptor and up regulates 5HT_2A_ receptor expression (**Figure 6B**).

We have also shown, using both genetic and pharmacological manipulations, that both serotonin and dopamine are able to suppress seizures in *repo*>*Dube3a* flies. In a previous study, we showed that glial overexpression of *Dube3a* causes seizures and synaptic impairments in Drosophila concomitant with down-regulation, at the protein level, of the Na^+^/K^+^ pump ATPα (17, 47). Serotonin modulates glial Na^+^/K^+^-ATPase activity in the rat cerebral cortex and hippocampus, through the 5-HT_1A_ receptor signalling (37). In addition, dopamine activates Na^+^/K^+^-ATPases and Mg^2+^-ATPases in the presence of the brain enriched soluble fraction of the rat cerebral cortex (48). Using behavioral and live imaging data assaying glial cell K^+^ content, we show here that both serotonin and dopamine suppress seizures by modulating Na^+^/K^+^-ATPase activity, despite the decrease in overall ATPα protein levels, possibly though the activation of ATPα via PKA phosphorylation. A massive influx of K^+^ ions into glia helps to restore the action potential to basal levels, and suppress seizures. Our new model proposes that serotonin and dopamine can reduce neuronal hyperexcitability by activation of PKA in glia, which in turn, activates the residual ATPα not degraded by Dube3a, and restores the basal levels of K^+^ ions in the synaptic cleft **(Figure 9).**

**Figure 9.**
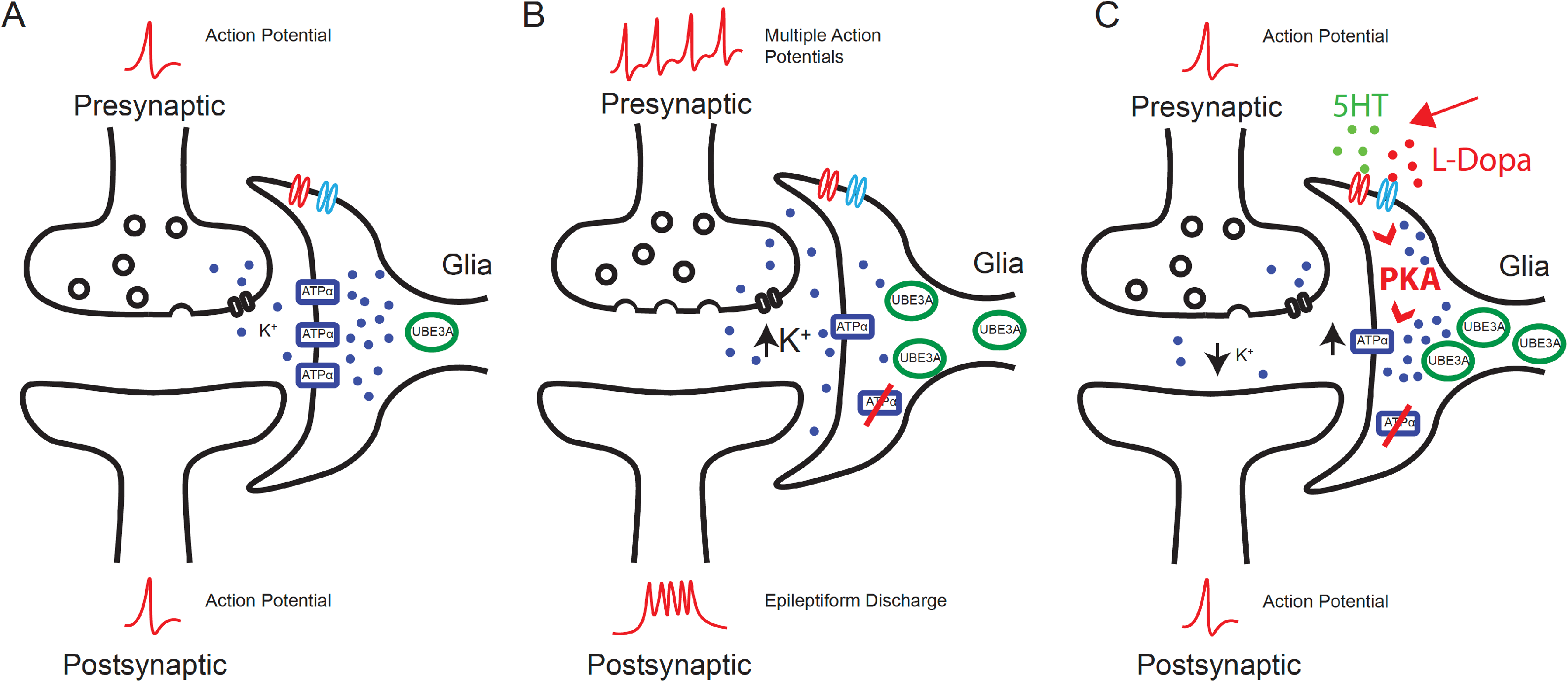
New model of ATPα modulation in Dup15q syndrome. A) Under normal conditions, the Na^+^/K^+^ exchanger ATPα pumps K^+^ ions from the synaptic cleft nearby glial support cells in order to maintain ionic homeostasis. B) When UBE3A levels are elevated in glia, the K^+^ concentration increases at the synapse because UBE3A can decrease ATPα levels in glia. The increase in K^+^ levels at the synapse leads to hyperpolarized neurons and pre-disposes these neurons to epileptiform activity. C) Here we have shown that a combination of 5HT and L-dopa can significantly suppress seizures in *repo*>*Dube3a* animals despite low levels of ATPα and increased Dube3a in glia. The primary method of this suppression is through the restoration of K^+^ imbalance. We have also documented a restoration of K^+^ levels in glial cells of the brain in the presence of 5HT + L-dopa. Here we propose a mechanism by which both 5HT and L-dopa can activate PKA, which in turn, activates residual ATPα within the glial membrane.

### Compounds Unrelated to Serotonin or Dopamine that Can Suppress Seizures

The compound screen we conducted also yielded some other potential seizure suppressors unrelated to serotonin or dopamine signaling. Even though we mainly focused on serotonin and dopamine for our mechanistic studies, these other hits are interesting as well since they regulate seizure activity via complementary mechanisms, which could prove useful in the future.

Pranoprofen, brand name Niflan, belongs to a class of medications called non-steroidal anti-inflammatory drugs (NSAID). Niflan can block the synthesis of prostaglandins by inhibiting cyclooxygenase, an enzyme converting arachidonic acid to cyclic endoperoxides, precursors of prostaglandins (49). Cytokines and prostaglandins are well-known inflammatory mediators in the brain, and their biosynthesis is enhanced following seizures (49). Dorzolamide hydrochloride is an inhibitor of carbonic anhydrases (CA), enzymes known to catalyze reversible hydration/dehydration of CO_2_/HCO_3_(−), respectively. CA inhibitors can reduce seizures through perturbation of the CO_2_ equilibrium and/or the inhibition of ion channels (50). Acetazolamide, a carbonic anhydrase inhibitor, is primarily used in combination therapy with other antiepileptic medications in both children and adults (50–52). Our findings support the use of dorzolamide hydrochloride as a potential anti-epileptic drug specifically for individuals with seizure caused by Dup15q syndrome.

Vital dyes like brilliant vital red (53) and methyl blue IV (54) have been proposed as anti-convulsive agents as far back as the 1930s. The prevailing hypothesis was that these dyes worked by rendering the blood–brain barrier impermeable to “convulsive toxins” in the systemic circulation (55). Recently, methylene blue has been found to be an anti-convulsant during self-sustaining status epilepticus (SSSE) induced by prolonged basolateral amygdala stimulation (BLA) in Wistar rats (56). Although speculative, it is possible that the contrast agent or dye Iopamidol, identified in our screen as a suppressor of seizures **(Figure 2**), might function in a similar manner.

### Advantages of using a Drosophila Model for Dup15q Seizure Suppressor Screening

Screening for compounds using a Drosophila model of a human disorder is not unprecedented. It has been clear for some time that genes across the fly genome show significant homology to human genes when associated with human disease etiology (57). Moreover, recent efforts have illustrated the use of Drosophila to quickly evaluate new disease associated variants *in vivo* (58–60) and compound screens have produced important new drugs for treating human conditions ranging from cancer to kidney stones (61–63), as well as nervous system conditions (64). In this study, for example, all of the genes in both serotonin and dopamine signaling, as well as the *UBE3A* gene, show high DIOPT scores of homology (65) to their human counterparts (**Table S3**). Functional studies in Drosophila support the idea that these basic neurotransmitter-signaling pathways are highly conserved from flies to humans as well (66–68). Finally, although several mouse models of Dup15q syndrome have been made, none of these mice show the spontaneous seizures found in patients, but our fly model does recapitulate the seizure sensitivity phenotype (17). Even if these mouse models did show a seizure phenotype, it would be practically impossible to screen 1280 compounds in a short period of time as we have done here.

One of the limitations of our Dup15q model, however, is that we are unable to do extensive behavior analysis, other than bang sensitivity, primarily because these flies are weak and very sensitive to seizure induction. Although we have shown recently that we can identify genes which affect ASD-like behaviors in flies (69) including repetitive behaviors (grooming), social communication (mating latency) and social spacing (social interaction) we are unable to utilize these behavior tests to explore the ASD aspects of Dup15q syndrome in our current fly model.

### Future Studies in Humans with Dup15q and Pharmacoresistant Seizures

Recently, our group identified an EEG biomarker in children with Dup15q syndrome, which has now been confirmed in other cohorts by high density EEG analysis (8, 70). This biomarker could potentially be used to assess the effects of treating children with Dup15q syndrome with 5-HT_1A_ agonists like vortoxitene in an off-label trial for seizure suppression in these children. Additionally, our other validated drugs have shown more effective seizure suppression in both male and female flies, than the routinely used medications for Dup15q patients (16) and refractory seizures (71), (72) (**Figure S5)**. Given the hint of success that treatment of non-syndromic epilepsy results in a decrease in seizure frequency when patients are given drugs for anxiety or depression (73), this study could significantly benefit the Dup15q community by providing a new, safe and a more effective method of decreasing seizure frequency.

## Acknowledgements

This work was initiated through a UTHSC CORNET Award to both G.P and L.T.R. and supported primary though NIH/NICHD R21HD091541 to L.T.R. K.A.H was supported by a predoctoral fellowship from the Dup15q Alliance. Stocks were obtained from the Bloomington Drosophila Stock Center (NIH P40OD018537) for this study. We also thank Yesenia Sanchez and Andrew Liess for their technical assistance with Drosophila stocks.

## Supplemental Information

**Table S1.** All validated compound descriptions.

**Table S2.** Drosophila stocks used from BDSC.

**Table S3.** DIOPT Homology Scores Between Drosophila and Human Genes.

**Figure S1.** Variance spread for males vs females in different solvents. Compounds in this study were dissolved in DMSO for the primary screen, but some compounds were soluble in either water or ethanol so the recovery times were assayed in both males and females in all three solvents. Males of the *repo*>*Dube3a* genotype showed a higher variance in all three solvents as compared to females. Red bars are SEM and black bar is the mean recovery time in that solvent. Error bars (red) are SEM. N > 40 animals per data point.

**Figure S2.** Drugs that failed to meet secondary screening go/no-go criteria. Error bars (red) are SEM. N > 40 animals per data point.

**Figure S3.** Drug Combinations that failed to significantly increase suppression over serotonin alone in both males and females. Error bars (red) are SEM. N>100 animals per data point.

**Figure S4.** Male fly results with 5-HT_1A_ agonists and antagonists. Error bars (red) are SEM. N>60 animals per data point.

**Figure S5**. Dup15q anti-seizure medications valproic acid and felbamate also suppress seizures in *repo*>*Dube3a* flies. Error bars (red) are SEM. N>50 animals per data point.

