## Supplemental Tables and Figures for "An unbiased drug screen for seizure suppressors in Dup15q syndrome reveals 5HT_1A_ and dopamine pathway activation as potential therapies"

### Supplemental Tables Figures

### Table S1

| Prestwick # | Chemical Name | Function | Predicted mode of action for seizure suppression | References |
|---|---|---|---|---|
| 1318 | **Pranoprofen** | Pranoprofen, brand name Niflan, belongs to a class of medications called non steroidal anti-inflammatory drugs (NSAID). It can be used to treat inflammation, keratitis, conjunctivitis and blepharitis after eye surgery. Its mechanism of action is the inhibition of inflammatory prostaglandin synthesis. | Cytokines and prostaglandins are well-known inflammatory mediators in the brain, and their biosynthesis is enhanced following seizures. Such inflammatory mediators could be therapeutic targets for the development of new antiepileptic drugs.It acts by blocking the synthesis of prostaglandins by inhibiting cyclooxygenase, which converts arachidonic acid to cyclic endoperoxides, precursors of prostaglandins. | Shimada et al., 2014 |
| 481 | **Serotonin hydrochloride** | Agents that elevate extracellular serotonin (5-HT) levels, such as 5 - Hydroxytryptophan and serotonin reuptake blockers, inhibit both focal and generalized seizures ; 2.Na+/K(+)-ATPase is activated also by 5-HT. | Several anti-epileptic drugs increase endogenous extracellular 5-HT concentration. 5-HT receptors are expressed in almost all networks involved in epilepsies. Currently, the role of at least 5-HT1A, 5-HT2C, 5-HT3 and 5-HT7 receptor subtypes in epileptogenesis and/or propagation has been described. Mutant mice lacking 5-HT1A or 5-HT2C receptors show increased seizure activity and/or lower threshold. | Bagdy et al., Journal of Neurochemistry, 2007 ; Brodie et al., Pharmacological Reviews, 2016; Chugani et al., Epilepsy Curr 2005; Hernandez Neurochem Intl 1992. |
| 871 | **Iopamidol** | Iopamidol is an organic iodine compound and used as a non-ionic water soluble radiographic contrast medium. | Unknown | |
| 1144 | **Mirtazapine** | Mirtazapine has a unique method of action by enhancing norepinephrine and serotonin neurotransmission by blocking the alpha-2 presynaptic adrenoceptors resulting in increased release of serotonin at the nerve terminals. | Mirtazapine actions at serotonin (5HT) synapses. : When presynaptic alpha 2 heteroreceptors are blocked by mirtazapine, 5HT is released with the potential to activate any 5HT receptors. However, because mirtazapine blocks 5HT2C, 5HT2A, and 5HT3 receptors, the increased serotonin release is directed largely to the 5HTlA receptor. | Schwasinger-Schmidt et al.Handb Exp Pharmacol. 2018; Kuhn et al. (2003) |
| 1116 | **Dorzolamide hydrochloride** | Dorzolamide is a carbonic anhydrase (CA) inhibitor. It is used in ophthalmic solutions (Trusopt) to lower intraocular pressure (IOP) in open-angle glaucoma and ocular hypertension. | Carbonic anhydrases are metalloenzymes that catalyze the reversible hydration/dehydration of CO(2)/HCO(3)(-), respectively. CA inhibitors can reduce seizures through perturbation of the CO(2) equilibrium and/or the inhibition of ion channels. | Aggarwal et al., Expert Opin Ther Pat. 2013 ; Millichap et al., Journal of pharmacology and experimental therapeutics, 1955. Acetazolamide, a carbonic anhydrase inhibitor,is primarily used in combination therapy with other antiepileptic medications in both children and adults. Reiss and Oles, Annals of Pharmaco Therapy, 1996 |
| 560 | **Prenylamine lactate** | Prenylamine has two primary molecular target in human, Calmodulin and Myosin light-chain kinase 2 found in skeletal and cardiac muscle. Pharmacologically, it decreases the sympathetic stimulation on cardiac muscle predominantly through partial depletion of catecholamine via competitive inhibition of reuptake by storage granule. | Depletes noradrnealine and dopamine from the reuptake by storage granules. Thereby increasing their concentration in the synaptic cleft. | |
| 66 | **Minaprine dihydrochloride** | Minaprine binds to serotonin type 2 receptors and to dopamine D1 and D2 type receptors. It also binds to the serotonin reuptake pump. Therefore, minaprine blocks the reuptake of both dopamine and serotonin. | minaprine appears to be a chemically and pharmacologically original antidepressant drug which activates both 5-HT- and DA-mediated transmission. | Biziere et al., Drugs Exp Clin Res. 1985 |
| 17 | **Levodopa** | The role of DA in epilepsy is still debated, although there is evidence of dopaminergic system involvement in certain animal models of epilepsy and in various forms of epilepsy in humans (Starr, 1996). In particular, alterations of subcortical dopaminergic pathways may be specifically related to the motor manifestations of certain types of seizures (Norden and Blumenfeld, 2002).In general, DA seems to exert an antiepileptic action, as demonstrated by the fact that the nonselective $D_1/D_2$ agonist apomorphine, with certain limitations, has anticonvulsant properties, whereas neuroleptic drugs that act as $D_1$ $D_2$ antagonists have predominantly proconvulsant actions. | There are two types of dopaminergic receptors, called the $D_1$ and the $D_2$. The former catalyzes the synthesis of cAMP, and the latter inhibits its synthesis. These reactions then regulate calcium and potassium channels in the postsynaptic membrane. | Brodie et al., Pharmacological Reviews, 2016 |

**Table S2**

| Genotype | Source | Stock Number |
|---|---|---|
| *Ddc* $^{DE1}$ | Bloomington Drosophila Stock Center | BDSC#3168 |
| *5HT1A* $^{D5kb}$/CyO | Bloomington Drosophila Stock Center | BDSC#27640 |
| *DAT* $^{Z21744}$ | Bloomington Drosophila Stock Center | BDSC#30867 |
| *Ddc* $^{27}$ *pr* $^{1}$/CyO | Bloomington Drosophila Stock Center | BDSC#3190 |
| *Ddc* $^{k02104}$ | Bloomington Drosophila Stock Center | BDSC#10508 |
| *ple* $^{4}$/TM3,*Sb* | Bloomington Drosophila Stock Center | BDSC#3279 |
| *Trh* $^{c01440}$ | Bloomington Drosophila Stock Center | BDSC#10531 |
| *5-HT2A* $^{C1644}$ | Bloomington Drosophila Stock Center | BDSC#4830 |
| *10XUAS-IVS-myr::tdTomato* | Bloomington Drosophila Stock Center | BDSC#32222 |
| *repo-GAL4 / TM3Sb* | Bloomington Drosophila Stock Center | BDSC#7415 |
| UAS-*ATPα* | FlyORF | F001458 |
| UAS *Dube3a* | Reiter et al., 2006, PMID: 16905559 | N/A |
| UAS human *UBE3A* | Reiter et al., 2006, PMID: 16905559 | N/A |
| *elav-GAL4* | Hugo Bellen (Baylor College of Medicine) | N/A |

**Table S3**

| Human Gene Name | Human Symbol | FlyBaseID | Fly Symbol | DIOPT Score | Rank | Best Score |
|---|---|---|---|---|---|---|
| *UBE3A* | *UBE3A* | FBgn0061469 | *Dube3a* | (14/18) | **high** | Yes |
| *Tyrosine Hydroxylase* | *TH* | FBgn0005626 | *ple* | (14/18) | **high** | Yes |
| *DDC* | *DDC* | FBgn0000422 | *Ddc* | (14/18) | **high** | Yes |
| *DAT* | *SLC6A3* | FBgn0034136 | *DAT* | (8/18) | moderate | Yes |
| *SERT* | *SLC6A4* | FBgn0010414 | *SerT* | (11/18) | **high** | Yes |
| *TPH1* | *TPH1* | FBgn0035187 | *Trh* | (13/18) | **high** | Yes |
| *5-HT1A* | *HTR1A* | FBgn0004168 | *5-HT1A* | (8/18) | **high** | Yes |
| *5-HT2A* | *HTR2A* | FBgn0087012 | *5-HT2A* | (5/18) | moderate | Yes |

Figure S1

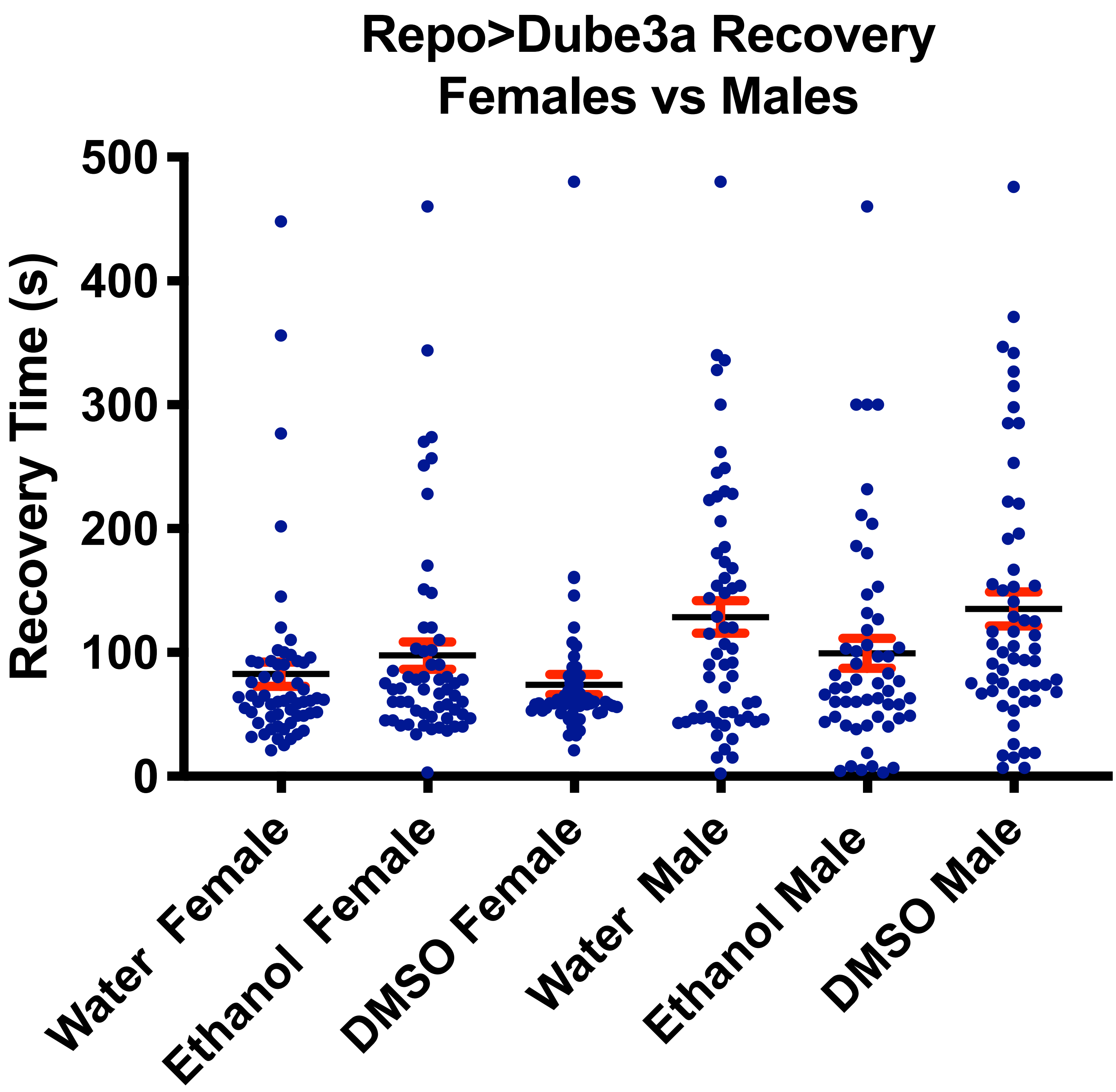

Repo>Dube3a Recovery
Females vs Males
Recovery Time (s)
500
400
300
200
100
0
Water Female
Ethanol Female
DMSO Female
Water Male
Ethanol Male
DMSO Male

Figure S2

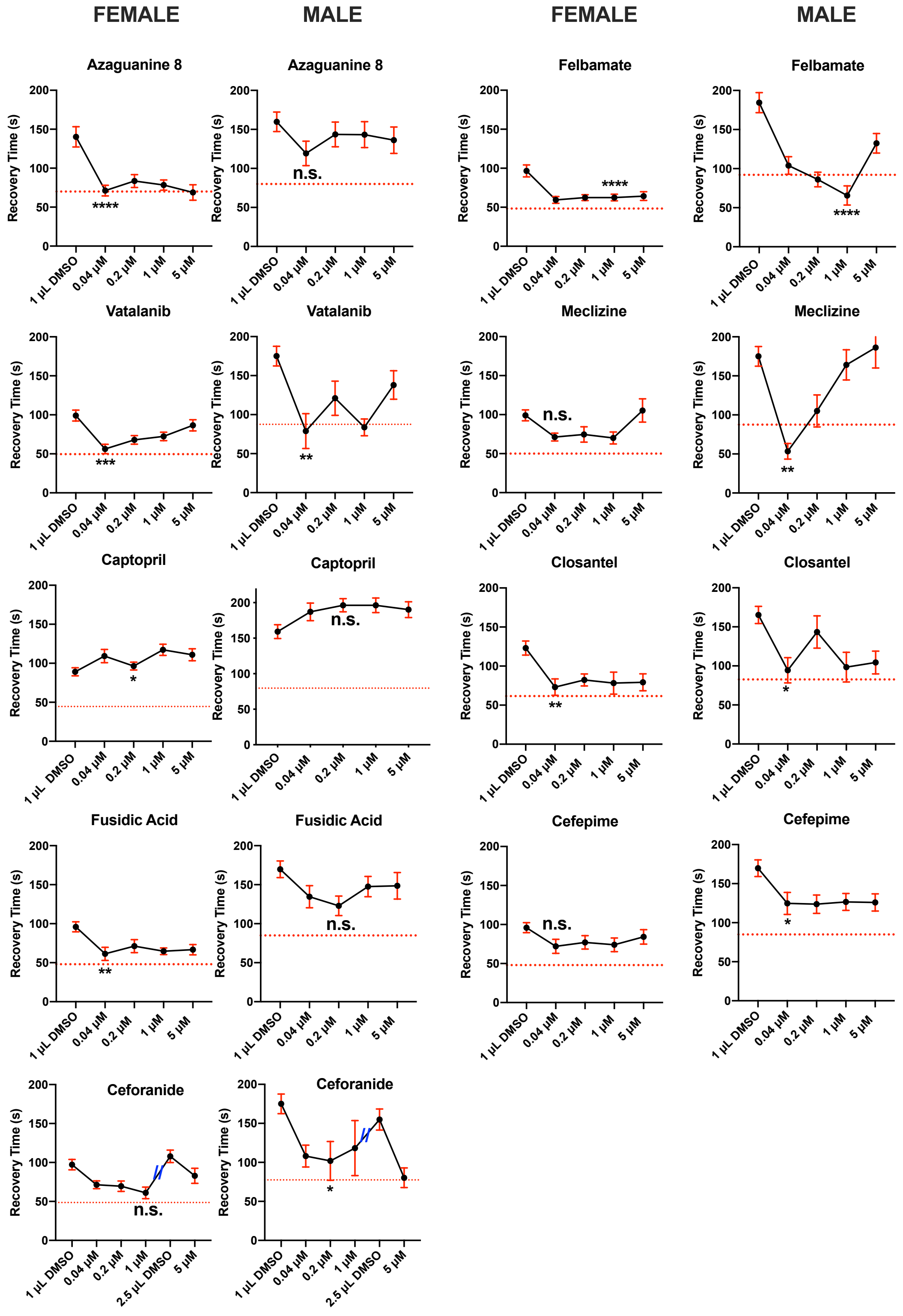
FEMALE
MALE
FEMALE
MALE
Azaguanine 8
Azaguanine 8
Felbamate
Felbamate
Vatalanib
Vatalanib
Meclizine
Meclizine
Captopril
Captopril
Closantel
Closantel
Fusidic Acid
Fusidic Acid
Cefepime
Cefepime
Ceforanide
Ceforanide
Recovery Time (s)
1 µL DMSO
0.04 µM
0.2 µM
1 µM
5 µM
2.5 µL DMSO
n.s.
****
***
**
*

**Figure S3**

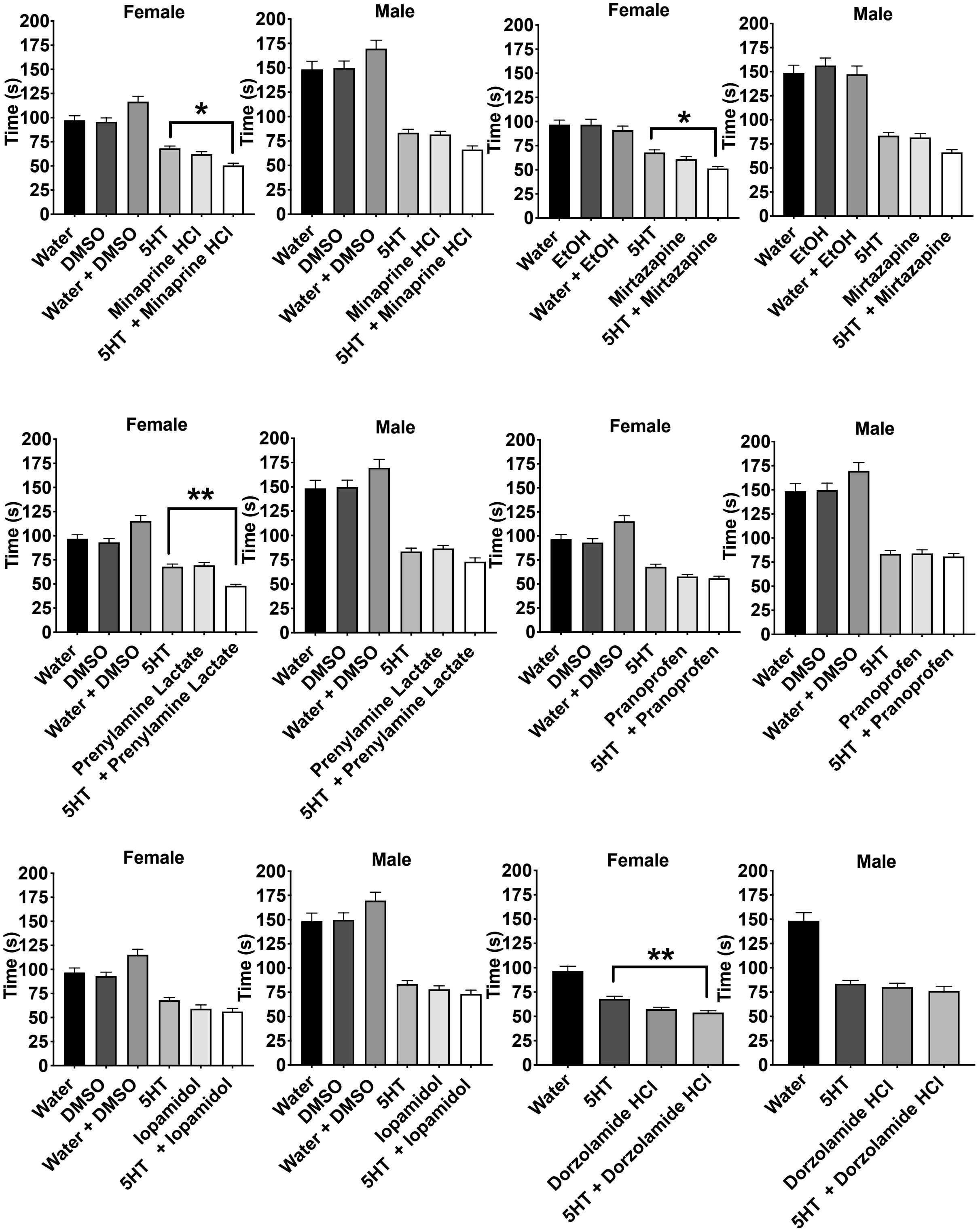
Female
Male
Time (s)
0
25
50
75
100
125
150
175
200
Water
DMSO
Water + DMSO
5HT
Minaprine HCl
5HT + Minaprine HCl
*
EtOH
Water + EtOH
Mirtazapine
5HT + Mirtazapine
Prenylamine Lactate
5HT + Prenylamine Lactate
**
Pranoprofen
5HT + Pranoprofen
Iopamidol
5HT + Iopamidol
Dorzolamide HCl
5HT + Dorzolamide HCl

Figure S4

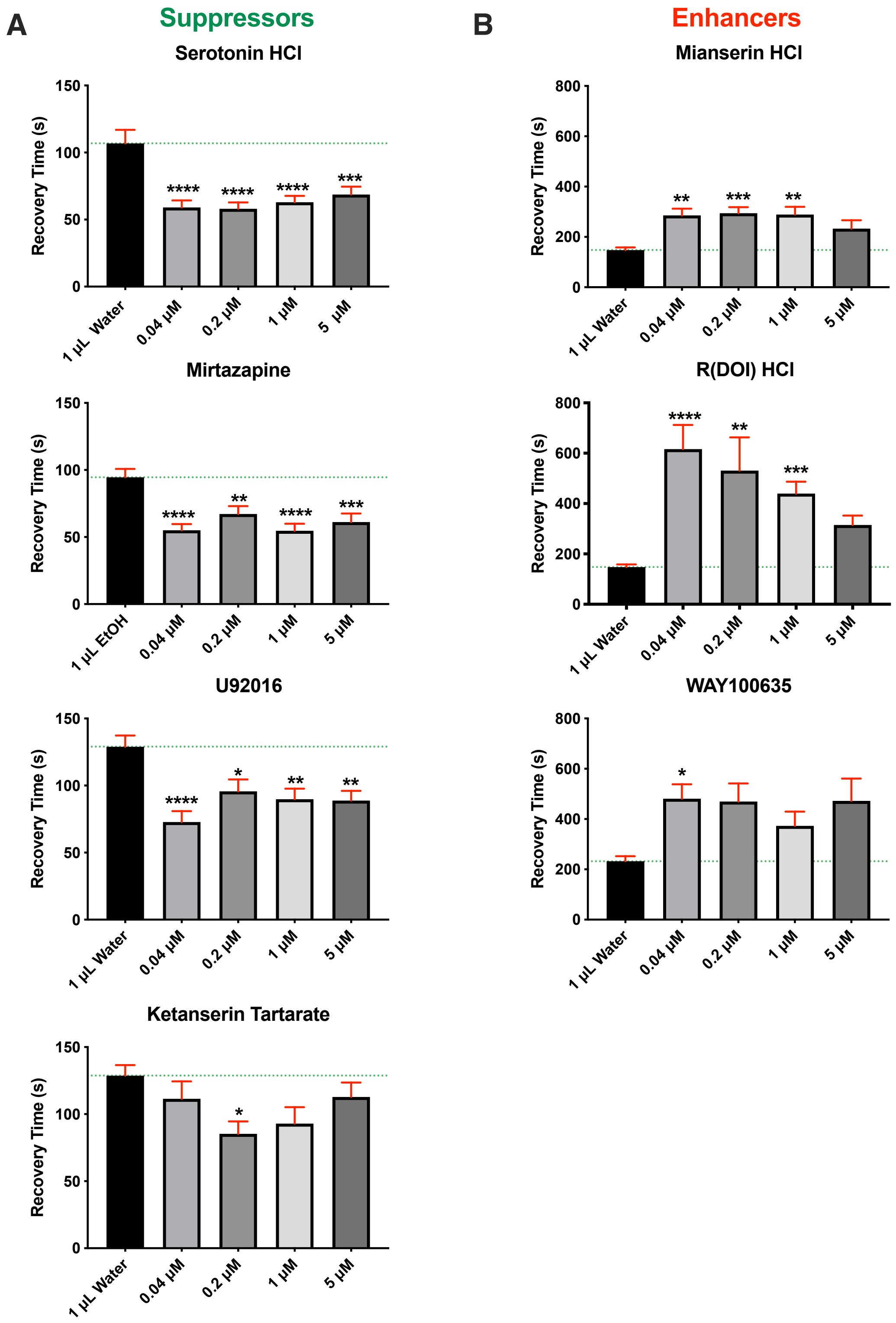
A
Suppressors
Serotonin HCl
Recovery Time (s)
150
100
50
0
****
****
****
***
1 µL Water
0.04 µM
0.2 µM
1 µM
5 µM
Mirtazapine
Recovery Time (s)
****
**
****
***
1 µL EtOH
0.04 µM
0.2 µM
1 µM
5 µM
U92016
Recovery Time (s)
****
*
**
**
1 µL Water
0.04 µM
0.2 µM
1 µM
5 µM
Ketanserin Tartarate
Recovery Time (s)
*
1 µL Water
0.04 µM
0.2 µM
1 µM
5 µM
B
Enhancers
Mianserin HCl
Recovery Time (s)
800
600
400
200
0
**
***
**
1 µL Water
0.04 µM
0.2 µM
1 µM
5 µM
R(DOI) HCl
Recovery Time (s)
****
**
***
1 µL Water
0.04 µM
0.2 µM
1 µM
5 µM
WAY100635
Recovery Time (s)
*
1 µL Water
0.04 µM
0.2 µM
1 µM
5 µM

Figure S5

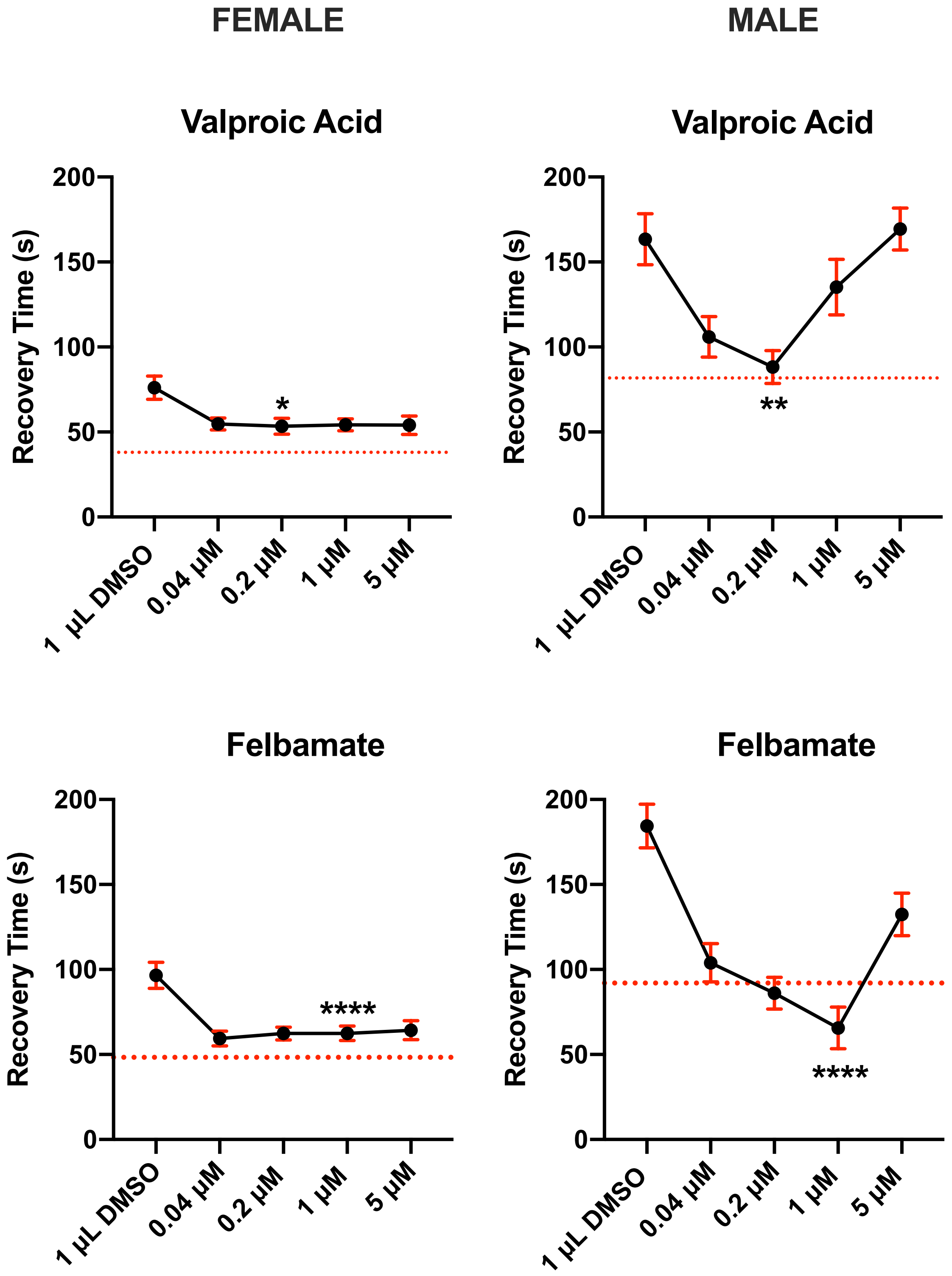
FEMALE
MALE
Valproic Acid
Valproic Acid
Felbamate
Felbamate
Recovery Time (s)
0
50
100
150
200
1 µL DMSO
0.04 µM
0.2 µM
1 µM
5 µM
*
**
****
****
